# Analog Signalling with ‘Digital’ Molecular Switches

**DOI:** 10.1101/256032

**Authors:** Stephen E Clarke

## Abstract

Molecular switches, such as the protein kinase CaMKII, play a fundamental role in cell signalling by decoding inputs into either high or low states of activity; because the high activation state can be turned on and persist after the input ceases, these switches have earned a reputation as ‘digital’. Although this on/off, binary perspective has been valuable for understanding long timescale synaptic plasticity, accumulating experimental evidence suggests that the CaMKII switch can also control plasticity on short timescales. To investigate this idea further, a non-autonomous, nonlinear ordinary differential equation, representative of a general bistable molecular switch, is analyzed. The results suggest that switch activity in regions surrounding either the high- or low-stable states of activation could act as a reliable analog signal, whose short timescale fluctuations relative to equilibrium track instantaneous input frequency. The model makes intriguing predictions and is validated against previous work demonstrating its suitability as a minimal representation of switch dynamics; in combination with existing experimental evidence, the theory suggests a multiplexed encoding of instantaneous frequency information over short timescales, with integration of total activity over long timescales.

**Author Summary:** Bistable molecular switches can decode cellular inputs into distinct high- or low-states of persistent enzymatic activity. Although this on-off, ‘digital’ perspective is valuable for long timescales, I suggest that short timescale fluctuations of switch activity around either stable state acts as an analog signal that reliably encodes instantaneous input frequency. A minimal model and theory make predictions about the molecular switch CaMKII, synaptic plasticity and burst detection.

## Introduction

Many cellular inputs lead to transient changes in cytosolic calcium (Ca^2^+) levels, generating temporally complex signals that reflect a wealth of information [1]. As such, cells express highly conserved molecular decoders capable of translating Ca^2^+ oscillations into downstream signalling events that affect diverse processes such as gene transcription, development and aging, neural network homeostasis and the synaptic plasticity that underlies learning and memory [2–9]. A celebrated example of a Ca^2^+ decoder is the protein kinase Ca^2^+/calmodulin (CaM)-dependent protein kinase II (CaMKII; Box 1), which can be suddenly driven by transient levels of cytosolic Ca^2^+ into either high or low states of switch-like activity. When stabilized through negative regulation by protein phosphatases, self-exciting (autophosphorylating) kinases such as CaMKII are an ideal component of signal amplification and have been previously likened to transistors on a computer chip, in that they may be turned on or off, presenting an ideal substrate for computation in cellular systems [10].

The classic CaMKII experiments of De Koninck and Schulman provided the first demonstration that a molecular switch can decode the frequency of periodic Ca^2^+ pulses into distinct, persistent levels of high enzymatic activation [11]. Although experimental evidence still largely lacks for whether this persistent activation occurs within functioning cells [12], there are recent indications that bistability does occur to some extent [13, 14] and that autophosphorylation is key to this process [15, 16]. Many modelling studies of CaMKII autophosphorylation dynamics capture the ability of the high activation state to persist beyond the original Ca^2^+ signal (known as hysteresis), which could potentially act over long timescales (seconds, minutes and longer) [17–19]. In these studies, the relationship between Ca^2^+ concentration and the state of the molecular switch are determined from simulations of detailed, parameterized systems of differential equations that are not readily amenable to deeper mathematical analysis; furthermore, these studies are restricted to periodic inputs and concerned with long timescale activation. In order to better understand frequency coding over short timescales (milliseconds to seconds) and its putative effect on synaptic plasticity (Box 1), this article analyzes a reduced description of molecular switch behaviour when subject to general aperiodic forcing and in the presence of noise, while further demonstrating the model’s compatibility with existing experimental and modeling results on long timescale CaMKII activation [11, 17]. As the study of cellular information processing shifts from individual transduction pathways, toward the emergent properties of complex signalling networks, simple mathematical models are becoming indispensable tools for both experimentalist and theoreticians alike [20, 21] by providing a trade-off between detailed performance and a reduced description that facilitates system-level studies. Much in the way that the leaky-integrate and fire model has benefited the study of spiking neurons [22, 23], the minimal switch model discussed in this paper will hopefully facilitate further study of complex kinase-phosphatase networks, while highlighting that a molecular ‘switch’ is more sophisticated than the name implies.

### Box 1: The bistable molecular switch CaMKII and synaptic plasticity

Accounting for approximately 1–2% of all brain protein, CaMKII is a central hub of cell signalling networks and can exert both pre- and post-synaptic control over information transmission in the central nervous system [24]. Once bound to the Ca^2^+-CaM complex, the kinase’s ability to cooperatively autophosphorylate can produce two distinct stable states: either high or low levels of enzymatic activation. Postsynaptically, after repetitive stimulation, the high activation state may persist after the Ca^2+^ signal subsides and can strengthen the connection between neurons, for example, the hippocampal CA3-CA1 synapses that support learning and memory [25]. However, it should be noted that the role of CaMKII autophosphorylation and bistability is not fully understood or accepted [12] and we are just beginning to gain better insight into the problem [15]. This paper proposes that CaMKII’s principal role is to meaningfully transmit information via its short term dynamics rather than store it permanently within levels of autonomously activated switch.

Presynaptically, CaMKII also modifies connection strength [26–28]. In weakly electric fish, the αCaMKII isoform produces presynaptic potentiation in a motion sensitive, excitatory sensory feedback pathway [28, 29]. The kinase also potentiates hippocampal CA3-CA1 synapses, as evidenced by knocking-out αCaMKII, which leads to reduced synaptic potentiation under paired pulse facilitation protocols when compared to the wild-type [30]. Through enzymatic phosphorylation of voltage gated Ca^2+^ channels and ryanodine receptors, αCaMKII can enhance Ca^2+^ entry and Ca^2+^-induced Ca^2+^ release in response to high frequency signals, potentially supporting hysteresis (Fig. 1) and driving synaptic release [31]. However, at the same CA3-CA1 synapses, post-tetanic potentiation protocols generate enhanced levels of potentiation in the same knock-out mice, illustrating that αCaMKII may also depress synaptic strength depending on the frequency and duration of the input [30]. Furthermore, αCaMKII has been shown to serve as a negative, activity-dependent regulator of neurotransmitter release probability at CA3-CA1 synapses [32]. This effect may be partially explained by the fact that CaMKII phosphorylates Ca^2+^-activated potassium channels that hyperpolarize the presynaptic terminal [33], decreasing the likelihood of Ca^2^+ entry and evoked neurotransmitter release. Intriguingly, αCaMKII also plays a non-enzymatic role in presynaptic CA3-CA1 plasticity by regulating the number of docked synaptic vesicles containing neurotransmitter [34]. In this case, decreased transmitter release could be explained by the fact that αCaMKII is acting as a sink for intracellular Ca^2^+, lowering the cytosolic levels that drive the machinery of synaptic vesicle fusion and influencing the size of the readily releasable vesicle pool [35, 36]. The size of the readily releasable pool is directly correlated with release probability at hippocampal synapses [37], supporting a putative role for αCaMKII in control of presynaptic plasticity parameters via Ca^2^+ and CaM buffering [32].

One of the most influential discoveries about CaMKII is its ability to decode the frequency of periodic Ca^2^+ pulses into distinct amounts of long lasting, autonomously activated kinase [11]. However, the interpretation of CaMKII as a frequency decoder has been criticized based on the fact that mean values of activity, evoked by different combinations of Ca^2^+ pulse size, duration and frequency, are ambiguously mapped into the same level of autonomously activated switch [38], which suggests that the switch is actually integrating the Ca^2^+ input over longer timescales. Alternatively, this article focuses on whether the concentration of activated switch acts as a reliable (analog) signal that reliably encodes frequency information over short timescales (sub-seconds), where Ca^2^+ pulse size and duration are far more stable [39]. The experimental evidence discussed above suggests that frequency coding by these ‘digital’ molecular switches is more sophisticated than previously thought and that fast fluctuations in presynaptic αCaMKII around either the stable high- or low-activation state can better represent instantaneous frequency information, and, hypothetically, translate it into bidirectional control of synaptic strength in real-time.

## Results

### A Bistable Switch Model

The following differential equation is an abstraction of a bistable molecular switch and was originally proposed as a model of genetic development by Lewis et al. [40]. This relatively simple model is a useful analytical tool to understand the general properties of bistable kinetic systems and captures the qualitative dynamics of more complicated models of CaMKII [19] (Fig. 1). Although the model interpretation and results presented here are centered on CaMKII and synaptic plasticity, the reader is encouraged to consider the broader implications for instantaneous frequency coding with other molecular switches, such as mitogen-activated protein kinases [7, 41].

**Figure 1.**
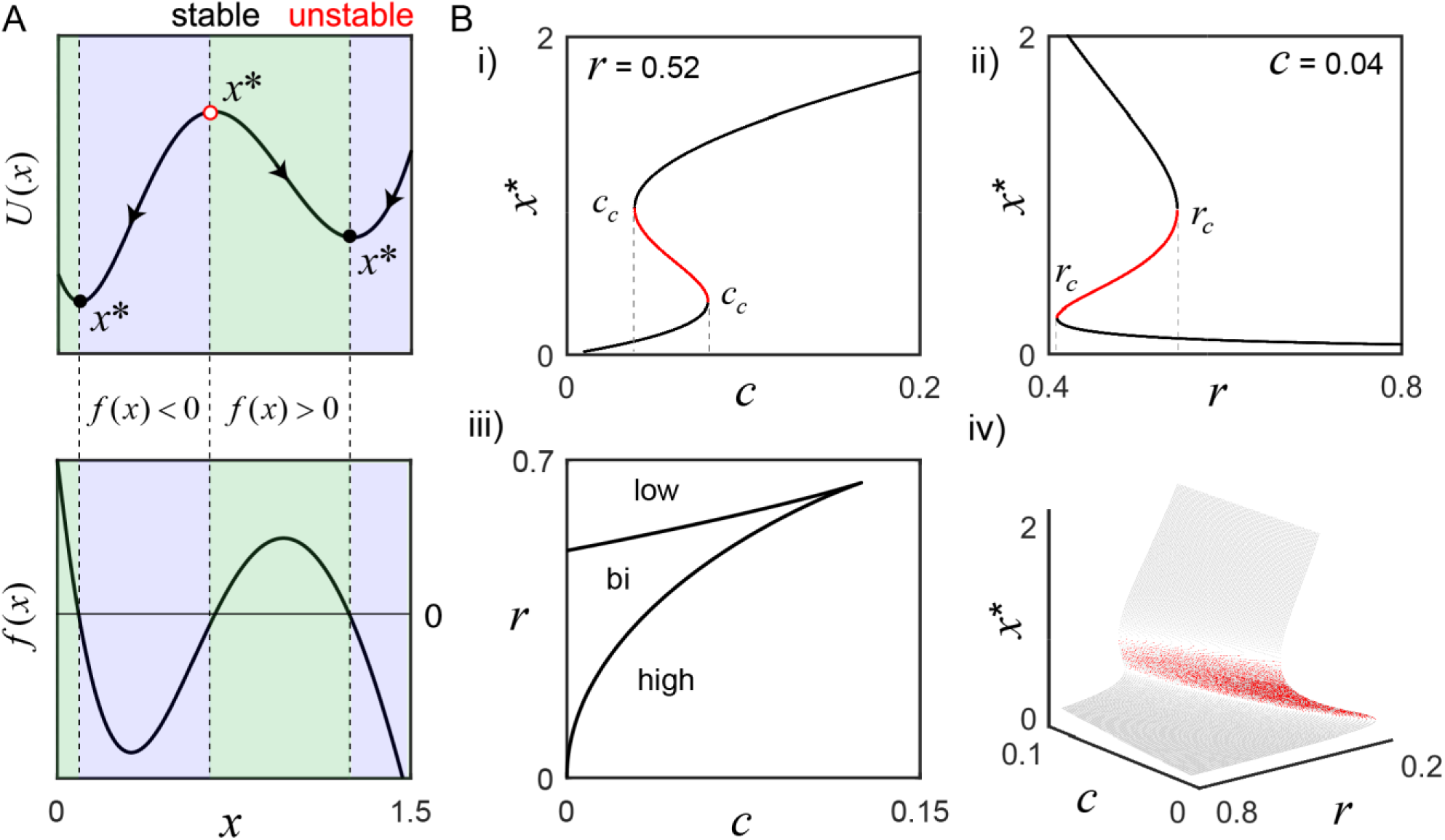
Activation states of the bistable molecular switch model. **A)** The model’s potential function, *U*(*x*), visually describes the tendency for solutions to settle around one of two equilibrium points (*x*\*), where the rate of change of switch activation, *f* (*x*), is 0 (parameters, *r* = 0.52 and *c* = 0.04). To the left of the stable equilibria (black circles), *f* (*x*) > 0 (green), and to the right, *f* (*x*) < 0 (blue), which forces perturbations to settle back into those states. Conversely, the sign of *f* (*x*) is reversed on both sides of the unstable equilibrium (red circle), such that tiny perturbations push the switch away, toward either stable state. **B)** As *r* or *c* change, *f* (*x*) changes and can result in the loss of bistability. **(i)** To illustrate, *r* is fixed as the input *c* is varied: small values only support low activation, but, as *c* grows, bistability emerges and eventually only the high activation state is supported when *c > c_c_* (rightmost). A defining feature of bistability is the hysteresis effect, where the same value of a parameter may evoke different states depending on the history of activity. For example, the high activation state still exists for *c* less than the rightmost *c_c_* and can only be lost when *c* falls below the leftmost *c_c_* value. **(ii)** *c* is fixed while the negative regulation parameter *r* is varied. For small *r*, only the high activation state exists. As *r* grows larger, the system becomes bistable and, eventually, only the low state exists after crossing *r_c_*. Panel **(iii)** shows a parametric plot of the critical values *c_c_* (*x*) and *r_c_* (*x*), and the bifurcation surface summarize the analysis completely **(iv)**.

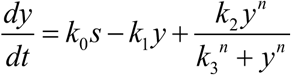

In this formulation, the level of activated CaMKII (*y*) is stimulated by the presence of Ca^2^+ bound to CaM, *s*, which will be studied as a function of time. For simplicity, it’s assumed that pulses of Ca^2^+ are bound upon cell entry, which is reasonable since CaM is found in large concentrations surrounding Ca^2^+ channels and has a strong affinity for Ca^2^+ [42]. Switch deactivation is directly proportional to the active CaMKII concentration at a rate *k*_1_, representing the activity of protein phosphatases. Finally, once activated, CaMKII has the ability to cooperatively bind Ca^2^+- CaM and autophosphorylate itself, which motivates the nonlinear, positive feedback term captured by the Hill equation, where *k*_2_ and *k*_3_ are the association and dissociation constants respectively. In addition to phosphorylation among the twelve subunits of a single CaMKII molecule, the ability to exchange active subunits between distinct CaMKII enzymes may connect this simple interpretation to a total, large pool of activated subunits distributed over multiple molecules [43]. Due to physiological constrains, *y*, *s*, *k*_0_, *k*_1_, *k*_2_, *k*_3_ ≥ 0. In the following, this specific equation will be referred to as the full kinetic model.

The full kinetic model of Lewis et al. has been previously applied to bistable genetic networks [40, 44, 45], transcriptional regulation [46, 47], mitogen-activated protein kinases [41], and incorporated into a larger phenomenological model of presynaptic plasticity [48]. Although insightful for their specific systems, these studies retain a large numbers of parameters that clutter analysis and obscure the generality of the results. Therefore, it is desirable to reduce the number of parameters and facilitate the following analysis by performing routine nondimensionalization. Let *y = x* · *k*_3_, 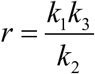, 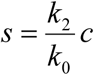 and 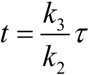, which, when substituted into the original equation and simplifying gives the reduced but dynamically equivalent form:

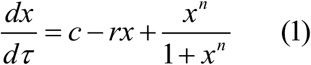

This article is interested in a time varying *c* ≡ *c*_0_ + *c_l_* (*τ*), where *c*_0_ reflects residual cytosolic Ca^2^+, whose slow dynamics are treated as fixed on the fast timescales over which the local Ca^2^+ signal *c_l_* (*τ*) fluctuates [49]. The timescale factor 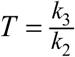, the quotient of the switch deactivation and activation parameters, will be reintroduced later in order to connect the switch dynamics to time in seconds and stimulation frequency in Hz. The parameter *r* represents the kinetics of CaMKII subunit dephosphorylation by protein phosphatases and scales with the factor *T*. Finally, for highly cooperative reactions, *n* = 2 is a reasonable approximation of the Hill function exponent [50] and a convention maintained by all of the studies listed above. The following bifurcation analysis is illustrated for *n* = 2, which allows for exact analytical solutions (Fig. 1 and Methods); however, the main results are then generalized to arbitrary *n* ∈ ℝ^+^, which is much more realistic and has important consequences for frequency coding.

### Stability and Bifurcation Analysis

Although interested in frequency-driven fluctuations over short timescales (Box 1), we must first examine the bistable, long timescale equilibrium behaviour of the model that defines the switch’s low and high activation states (Equation 1; Fig. 1). An important reason for reducing the number of model parameters above is to simplify the analysis of all the possible system behaviours as a function of only a few parameter values. Having selected *n* = 2, we now only need to consider the effect of varying *r* and *c*; depending on their values, we may have one, two or three equilibrium points (*x*\*), where the rate of change of the switch 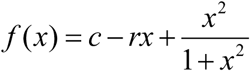 is equal to zero. For example, consider the values *r =* 0.52 and *c =* 0.04 that support bistability: there are three fixed points, two of which are stable, as illustrated by the switch’s potential function *U*(*x*) = −*∫ f* (*x*)*dx* (Fig. 1A). As *r* and *c* change, saddle node bifurcations can occur, resulting in the presence of only the high or low activation state. The corresponding bifurcation diagrams are displayed in Figure 1B; their derivation is found in the Methods section.

A key feature of bistability is the hysteresis effect, where the same value of a parameter may evoke different states depending on the history of activity. For example, as the Ca^2^+ signal *c* increases, *x*\* grows larger until crossing the rightmost *c_c_*, where a saddle node bifurcation occurs and the switch jumps up to the high activation state, as the low state disappears (Fig. 1Bi). Now, as *c* decreases back into the bistable range, the high activation state is preserved, and only lost when *c* crosses below the leftmost value of *c_c_*. This history dependent behaviour is presumably central to sustained CaMKII activity on the order of seconds [28] (Box 1). A similar phenomenon occurs for the negative regulation parameter *r* (Fig. 1Bii). The values of *r_c_* and *c_c_* are plotted parametrically as a function of the active switch in the bifurcation curves (Fig. 1Biii). The bifurcation surface summarizes this information completely (Fig. 1Biv).

### Existence of Solutions Around Stable Equilibria

To date, studies of Lewis et al.’s full kinetic model have been restricted to static input and periodic forcing. It is of principal interest to characterize the model behaviour in response to aperiodic forcing, in order to gain a more general, physiologically realistic understanding of frequency coding with molecular switches. In addition to potentially encoding frequency information into stable levels of activated switch for many seconds presynaptically [28], or minutes postsynaptically [11, 25], what about frequency coding on the order of milliseconds to seconds, which is associated with brief sequences of action potential-evoked Ca^2^+ inputs? In a neighbourhood surrounding a stable activation state (a sub-state region of state space), is there a unique solution for a given time varying input signal? This question is not trivial, since small changes in the initial conditions of a nonlinear system (i.e., past switch activity) may generate drastically different behaviours. Understanding the relationship that determines whether solutions converge or diverge around a given steady state could provide valuable insight into the properties of bistable molecular switches.

In the following section, we now reintroduce the scale factor *T*, since we are interested in studying frequency in Hz and time (*t*) in seconds. As such, Equation 1 becomes

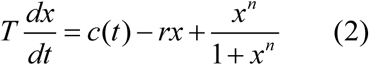

First, to establish the existence of solutions around the high and low switch states, consider Equation 2 and note that *f* explicitly depends on the time-varying forcing term, *c*(*t*) = *c*_0_ + *c_l_* (*t*). The phosphatase activity *r* that can counteract the switch phosphorylation is treated as fixed. The function *f*(*t*, *x*(*t*)) is assumed to be Lipschitz continuous and well-defined within intervals of state space, *y*_−_ ≤ *x*(*t*) ≤ *y*_+_ satisfying the conditions *f*(*t, y*_+_) > 0 and *f*(*t*, *y*_+_) < 0 for all *t* ∈ ℝ^+^ (recall Fig. 1A), which traps solutions within these boundaries unless the signal itself triggers a transition. For any given point in time, there exist boundaries (*y*_−_, *y*_+_) determined by the parameters *r*, *c*_0_ and the input *c_l_* (*t*); we refer to values of the activated switch falling within these trapping regions as sub-state solutions, that is, fast timescale fluctuations that occur around either the high or low stable activation states [25].

For (*c*, *r*) corresponding to the bistable region of parameter space (Fig. 1Biii), there exist two intervals, *x*(*t*) ∈ (*y_l_*_−_, *y_l_*_+_) and *x*(*t*) ∈ (*y_h_*_−_, *y_h_*_+_), each surrounding one of the stable equilibrium points (*x*\*).

Now, we wish to locate values for the low state (*y_l_*_−_, *y_l_*_+_) and high state (*y_h_*_−_, *y_h_*_+_), where the existence of local time-varying solutions can be established. This problem is intimately linked to bifurcation, since *y_l+_* and *y_h−_* depend on the values of *c* and *r*. The choice of a lower bound for the interval that exists around the low activation state is *y_l−_* = 0, since the physiological restriction *c*(*t*) ≥ 0 implies *f*(*t*, 0) > 0 for all *t* ∈ ℝ, ignoring the boring degenerate case of *c*(*t*) = *x*(*t*) = 0. The upper bound of the lower strip, *y_l_*_+_, can be chosen as a value 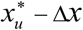, left of the unstable equilibrium 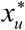 where 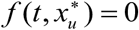, such that 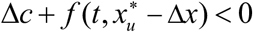; this condition ensures that the system is not trivially displaced into the high activation state by a single Ca^2^+ pulse with amplitude Δ*c*. For the high concentration strip (*y_h_*_−_, *y_h+_*), the lower bound *y_h_*_−_ is chosen as a value of *x* infinitesimally greater than 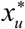, that is, 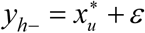 for *ε →* 0. Since we have restricted *r* and *c > c_c_* (leftmost; Fig. 1B) to the bistable range, we know that *f*(*t*, *y_h_*_−_) > 0. For the upper bound of the high activation strip, it is enough to note that for 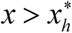, *f*(*t*, *x*(*t*)) < 0 and, since we wish to maximize the width of the strip, we take *x* arbitrarily large, denoting this value by *y_h+_ = x*_∞_. During stimulation, if (*c*, *r*) drifts out of the bistable region of parameter space, a saddle node bifurcation occurs and only one interval exists; in this case, the bounds simply span the state space, *y*_−_*=* 0 and *y*_+_ = *x*_∞_.

By invoking the Cauchy-Peano theorem, we guarantee the existence of at least one sub-state solution for every initial condition found within the interval regions defined above, since the conditions on the sign of the derivative *f*(*t*, *x*(*t*)) define trapping regions. However, this theorem says nothing about whether solutions starting at different initial conditions will converge to a unique, stimulus-driven response that tracks changes in the Ca^2^+ signal.

### Uniqueness of Sub-state Solutions

As motivation for the following results, Figure 2A shows an example switch response to an 8 Hz Poisson pulse sequence, which is convolved with an alpha function filter (30 ms, Methods), then normalized to the signal’s maximum and scaled by Δ*c* = 0.5 to create an example input signal, which the switch tracks closely. Note, in this simulation, the alpha-function kernel was specifically chosen to be 30 ms based on literature values for the time course of local synaptic Ca^2^+ signals [51–53]. Due to our interest in the fast timescales associated with short sequences of input pulses (100s of milliseconds), we assume that the Ca^2^+ pulse size is fixed on this timescale, which is a reasonable approximation for hippocampal spiking frequencies less than 15 Hz [39]. This distinction between short and long timescales provides a hypothetical means for the system to be less sensitive to variations in the Ca^2+^ pulse size and the resulting frequency-intensity coding ambiguity ([19, 38]; see Box 1). This could allow for more accurate representations of instantaneous frequency over short time periods, compared to long timescale frequency coding where input history, as well as additional adaptive and homeostatic processes may substantially adjust Ca^2^+ signalling.

**Figure 2.**
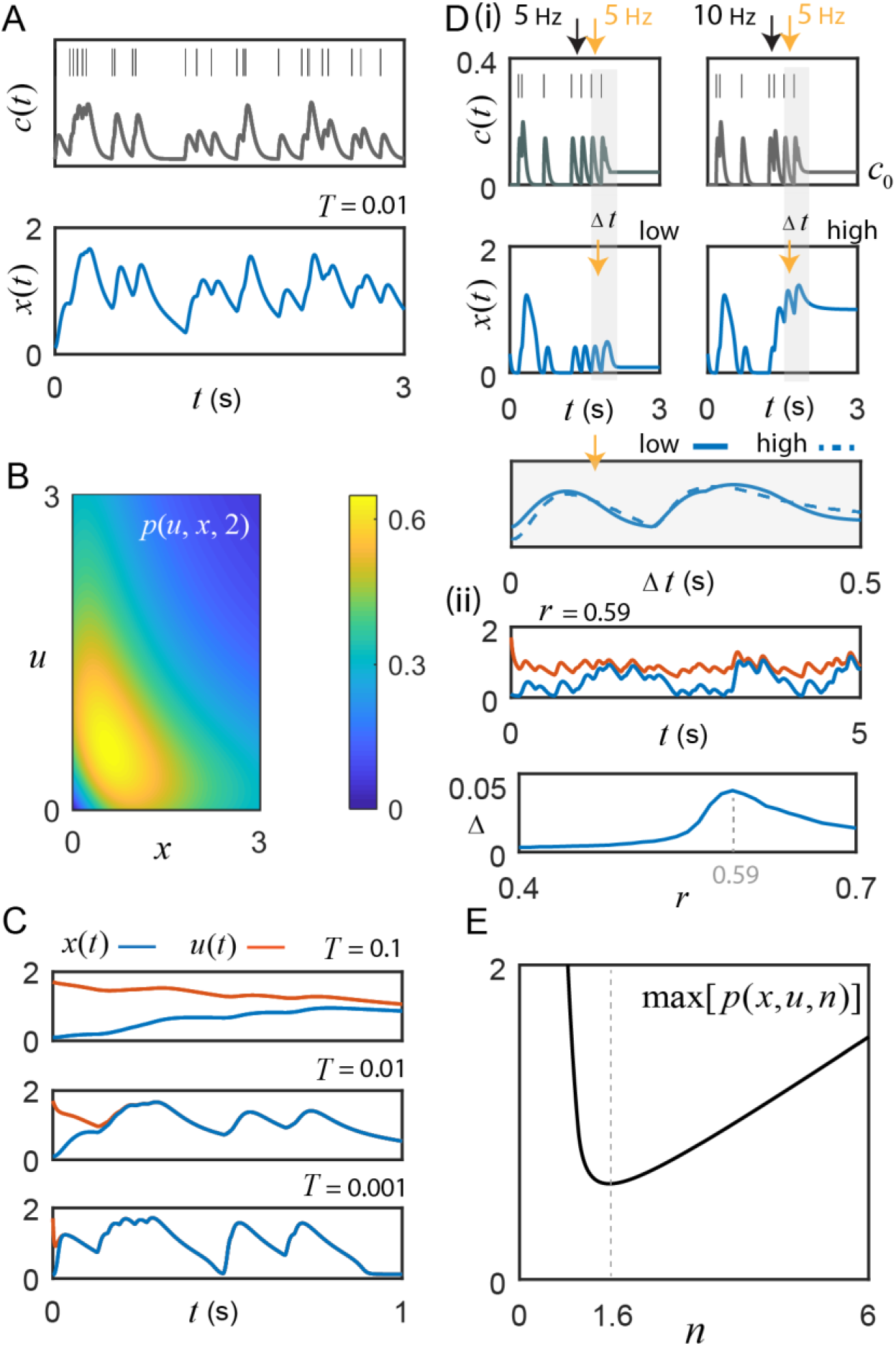
Switch activity fluctuates with instantaneous input frequency. **A)** Motivating example: switch response to an 8 Hz Poisson sequence of input pulses, convolved with an alpha function kernel to create a signal, *c*(*t*) = *c*_0_ + *c_l_*(*t*). The switch’s fluctuations track changes in the input frequency (*n* =1.6, *r* = 0.61, *c*_0_ = 0.04, and *T = 0*.01) **B)** The example function *p*(*u*, *x*, 2) from the uniqueness proof achieves a maximum of 0.65; *r* must exceed this value to guarantee absolute convergence of the switch to a unique frequency-driven solution. **C)** Initial conditions: *u*(0) =1.7 and *x*(0) = 0.1. The value of *T* affects time-to-convergence between solutions and frequency filtering. From empirical studies, *T* ≤ 0.01 [54]. **D) (i)** For *r* = 0.54 < *r_c_*, sufficiently high frequency Ca^2^+ pulses (bursts) cause transitions from the low to high state (illustrated for *n* — 2). By adjusting *c*_0_ to take advantage of hysteresis, the cell can control whether or not it is sensitive to these burst-induced up states. The first two pulses (<10 Hz), where *c*_0_ = 0, do not result in hysteresis, whereas the high frequency 10 Hz inter-pulse interval (right panel black arrow) with *c*_0_ augmented to 0.04 does; note that neither static value can generate the upstate alone without sufficient input (e.g., 5 Hz, left panel black arrow). The switch response differs during the transition between low and high states, but once settled around a given state gives good agreement (gray shading; the two example curves are compared by choosing an offset of 0.92 that minimizes the Euclidean distance between them). **(ii)** *Top* Simulation for *r* = 0.59 and *n* = 2, where *x*(*t*) has *c*_0_ = 0.02 and thus cannot support bistability, versus *u* (*t*) with *c*_0_ = 0.04, which, when driven by input, traps the solutions around the high activation state through hysteresis. Under these conditions, convergence cannot occur. *Bottom* The absolute value of the difference between the relative changes in *u* (*t*) and *x*(*t*) induced respectively by the common input frequencies (Δ, see Results for details) plotted as a function of *r*; the maximum discrepancy of 0.046 is found at *r* = 0.59 and represents a small fraction of the total activated switch concentration. **E)** In general, the exponent *n* ≠ 2 in real biological systems. Interestingly, *n* = 1.55 is a minimum for the maximum value of the class of functions *p*(*u*, *x*, *n*)in the uniqueness proof. This is remarkably close to the empirical best-fit value of 1.6 reported by De Koninck and Schulman for presynaptic αCaMKII [11].

We now establish the stability and uniqueness of solutions for distinct initial conditions within a given interval of state space. Consider a general interval (*y*_−_, *y*_+_), where *x*(*t*) is a solution to Equation 2 with initial condition *x*_0_ ∈ (*y*_−_, *y*_+_). Assume there is another solution, *u*(*t*), with a different initial condition *u*_0_ ∈ (*y*_−_, *y*_+_). Writing *z*(*t*) = | *u*(*t*) − *x*(*t*) | and first assuming *n* is a positive integer, we see that

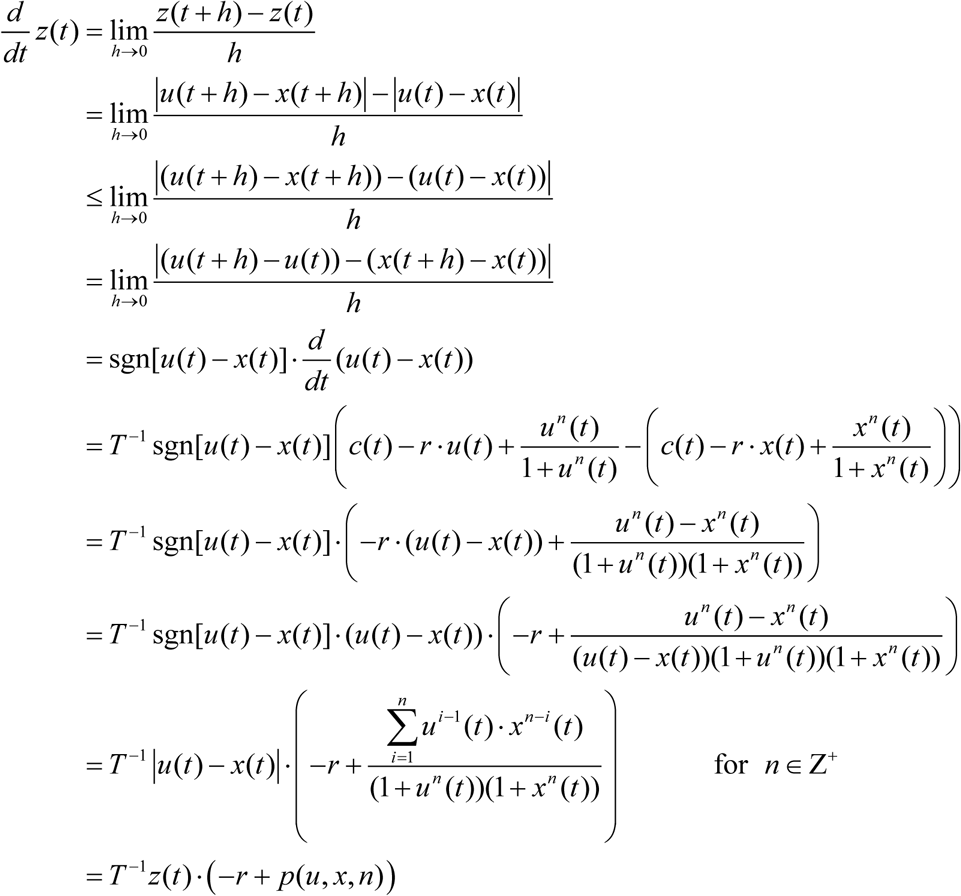

The expression *p*(*u*, *x*, *n*) achieves maximal values at intermediate switch levels that separate the low and high states of activation. Now, consider *p*(*u*, *x*, *n*) for the special case of *n* = 2, used in the bifurcation analysis; in this case, 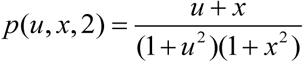, which is plotted in Figure 2B. Setting the partial derivatives of the function to zero and solving for *u* and *x*, yields a critical point: 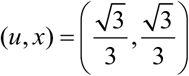. Substituting this into *p* gives a global maximum of 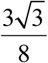. Since 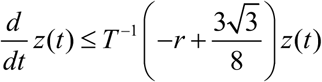 for all *t*, we can apply Grönwall’s inequality, which gives us the following:

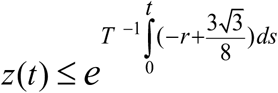

Substituting the expression for *z*(*t*) and solving this integral exponent yields,

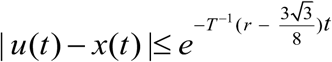
and, as *t* →∞, we have

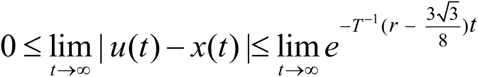

For 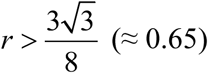, we obtain

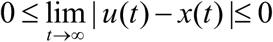

By the squeeze theorem we conclude that | *u*(*t*) − *x*(*t*) |→ 0 as *t* → ∞. Therefore, a unique frequency-driven solution exists and is independent of the initial conditions within the bounded interval. The time taken to converge to the unique solution is inversely proportional to 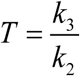 (Fig. 2C). The parameter value *T* = 0.01 seconds was chosen here for our specific example switch, CaMKII, whose dissociation constant (*k*_3_) has been experimentally determined to be at least 100-fold smaller than the activation constant (*k*_2_) that governs the rate of autophosphorylation [54]. Unlike the larger value of *T* = 0.1 seconds, *T* = 0.01 permits quick convergence and reliable encoding for the action potential frequencies characteristic of hippocampal CA3-CA1 synaptic input (approximately 1–15 Hz) [55]. Smaller values of *T* permit rapid convergence and more accurate frequency coding, but may become overly sensitive to temporary lulls in activity when *c* briefly drops below the leftmost critical value *c_c_* (recall Fig. 1Bi).

It should be noted that *r* > 0.65 is an absolute guarantee of convergence to a unique frequency driven solution; but, from the bifurcation analysis (Fig. 1Biii; Methods), we know that bistability does not exist for this value of *r*. However, in general, only *−r + p*(*u, x*, 2) < 0 is required, which, for low and high concentrations of activated switch, is obtained at smaller values of *r* that do support bistability. In fact, *p*(*u*, *x*, *n*) only exceeds the *r* value briefly during state transitions as it moves through the unstable equilibrium. Although a unique encoding of sub-state solutions can still exist for smaller *r* values around either the high or low state, convergence about the low activation state is now vulnerable to perturbation by short Ca^2^+ inter-pulse intervals, thus acting as a high frequency event (burst) detector through induction of high switch activation. For example, experiments show that high frequency hippocampal activity (>15 Hz) causes successive Ca^2^+ pulses to accumulate [39], which could effectively boost *c*_0_ and serve to promote burst detection via maintenance of hysteresis and the high activation state (Fig. 2Di). In theory, this dynamic burst threshold (the separatrix) is sensitive to recent levels of activation, and could be purposefully modulated by the cell through dynamic regulation of the parameters *r* and *c*_0_ [56]. To restore the low state, the cell simply needs to adjust *c*_0_ to fall below the leftmost critical value *c*_c_. The bottom panel of Fig. 2Di illustrates that fluctuations around the high- and the low-stable states still yield nice agreement in their response to a given input frequency. Of course, during the state transition itself, the switch response can differ largely but once solutions are settled around their respective stable states the model appears to give good agreement (gray shading; the two example curves are compared by choosing an offset of 0.92 that minimizes the Euclidean distance between them).

When bistability is supported, the model response cannot always converge to an absolute level of activated switch, as illustrated in the top panel of Fig. 2Dii; however, the fluctuations about the distinct stable states appear to be similar, as in Fig. 2Di. To examine this idea further, repeated simulations of the model were performed, where *x*(*t*) has an associated residual Ca^2^+ level of *c*_0_ = 0.02 and thus does not support bistability, versus *u* (*t*) with *c*_0_ = 0.04, which can trap the solution around the high activation state through hysteresis (Fig. 2Dii, top). As was the case in Fig. 2C, the same random spike sequences are used for *x*(*t*) and *u* (*t*) on each trial. For each inter-pulse interval of the repeated simulations, the change in the level of activated switch was computed as the difference between the switch activity sampled at the time of an input pulse and the subsequent maximum switch response that occurred before the next pulse. For each successive, shared inter-pulse interval, these differences, Δ*x* and Δ*u*, were determined separately for *x*(*t*) and *u*(*t*), then subtracted from each other for each 100 second trial, containing an average of 797 pulse intervals (8 Hz Poisson process). This was repeated 10 times for each parameter set and the composite mean of the absolute value of the difference between the change in the two solutions, Δ = |Δ*u* − Δ*x*|, was determined as a function of *r* (Fig. 2Dii, bottom). The maximum discrepancy between Δ*x* and Δ*u*, 0.046, occurs at *r* = 0.59 (used for Fig. 2Dii, top) and is at least an order of magnitude less than typical values achieved in the low activation state. These results suggest that the relative change in switch activation about a stable state is generally quite consistent.

Realistically, the Hill function exponent *n* need not be restricted to integer values, which is unlikely in real biological systems. Thus, in the above proof, the expression *p*(*u*, *x*, *n*) is now left as 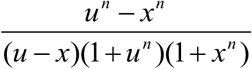 for *n ∈* ℝ^+^, since there is no longer a closed form expression for the factorization of the numerator by the term *u − x*. The function *p*(*u*, *x*, *n*) has critical points at *u = x*, which occur at an apparent discontinuity due to the factor *u − x* in the denominator. However, assessing the limit as the difference between *x* and *u* becomes infinitesimally small, making the change of variable *u = × + h* as *h* → 0, and recognizing the limit definition of the power rule for differentiation, yields an expression for the maximum of *p*(*u*, *x*, *n*) for all *u, x, n* ∈ ℝ^+^:

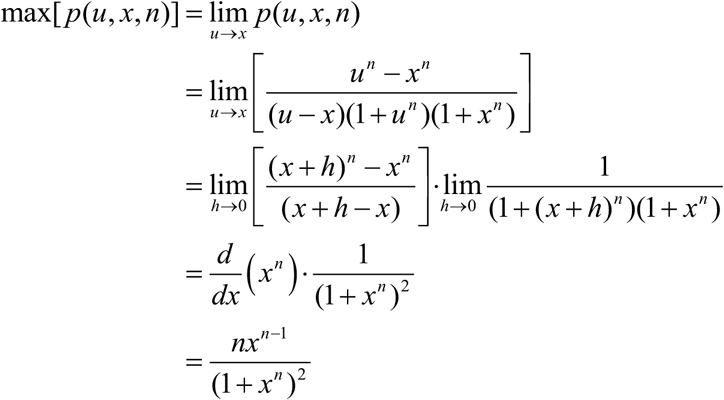

For each value of the exponent *n*, the global maximum of this expression is determined for all *x* ∈ ℝ^+^, and plotted (Fig. 2E). Ignoring the highly uncooperative reaction exponents of *n <* 0.012, the global minimum of the class of functions *p*(*u*, *x*, *n*) is found at *n* = 1.55. Fascinatingly, the empirical αCaMKII data reported by De Koninck and Schulman [11, 17] was fit by a Hill function with an exponent of 1.6. This intriguing match between their experiment and the model’s theory suggests that αCaMKII’s activation function may operate with this particular exponent as it provides the minimum level of negative regulation *r* required to maintain absolute convergence of unique input driven switch activity in the low activation state, even for intermediate levels of the switch response occurring just left of the unstable equilibrium (Fig. 1A), where *r* must be much stronger to guarantee unique solutions (Fig. 2B). As we will see in the following section, the value of *n* = 1.6 has additional benefits for amplifying the frequency response of weak calcium fluctuations in the presence of noise.

### Molecular Switches and Stochastic Resonance

If Equation 2 is to capture actual molecular switch behaviour *in vivo*, then we must understand frequency coding in the presence of biological noise. Given our interest in synaptic information transfer, it is natural to ask whether noise can improve the switch’s frequency coding ability through stochastic resonance and how different combinations of our main parameters (for example the value of *n*) could potentially affect this phenomena. In particular, does the value *n* = 1.6 confer benefits for frequency coding? The results presented in this section are generated by Equation 2 with additive Ornstein-Uhlenbeck noise, *η*(*t*), which evolves according to the stochastic differential equation

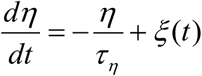
where *ξ*(*t*) is bounded Gaussian noise, *N*(0,1), whose amplitude is scaled by a parameter *σ*. The simplest interpretation is that there is some weak noise in the Ca^2^+ signal amplitude, which might arise from stochastic channel dynamics. The choice of the time constant *τ_n_* is based on previous studies of noisy microdomain Ca^2+^ fluctuations, where an upper bound for the autocorrelation time was determined to be approximately 10 ms [57, 58]. This choice has the added benefit of matching our switch time constant *T*, should we instead assume the noise is inherent to switch activation, as well as matching a typical value for the membrane time constant of spiking neurons, whose noisy membrane potential fluctuations might influence the activity timescales of voltage-gated Ca^2^+ channels.

Figure 3A shows the power spectrum (*P_c_*) of a weak sinusoidal Ca^2^+ oscillation, *c = c*_0_ *+α* sin(2*πφt*), where *c*_0_ = 0.04, *α* = 0.02 and *φ* = 2 Hz, which was selected based on the mean action potential frequency associated with the CA3 and CA1 regions of the hippocampus [59]. As expected, the noisy switch oscillates at the frequency *p*, reflected in its power spectrum (*P_x_*). Very recently, the full kinetic model of Lewis et al., studied under the context of genetic regulation with *n* = 2, has been shown to produce the stochastic resonance effect [47], which is confirmed here for the dimensionally reduced model (Equation 2; Fig. 3B). As *σ* increases from 0, frequency transfer, measured as the ratio of the switch power to signal power at *φ*, dips slightly and then improves dramatically, achieving a maximum at 0.29, followed by a quick decrease as the noise becomes dominant. When changing the exponent from *n* = 2 to *n* = 1.6, this spectral amplification becomes significantly larger, further suggesting that presynaptic αCaMKII functions as an important frequency decoder and that the exponent *n* = 1.6 may have evolved to fulfill this purpose. The reader should note that, for fair comparisons sake, *r* = 0.65 and *r* = 0.61 were selected respectively for *n* = 2 and *n* = 1.6 based on values obtained from Fig. 2E, but this effect is qualitatively robust to changes in *r* and *φ*. Setting *n* = 1.6 also shifts the optimal noise strength to a substantially lower value, 0.09, which has the putative benefit of harnessing stochastic resonance and enhanced frequency representations for low intensity Ca^2+^ signal noise.

**Figure 3.**
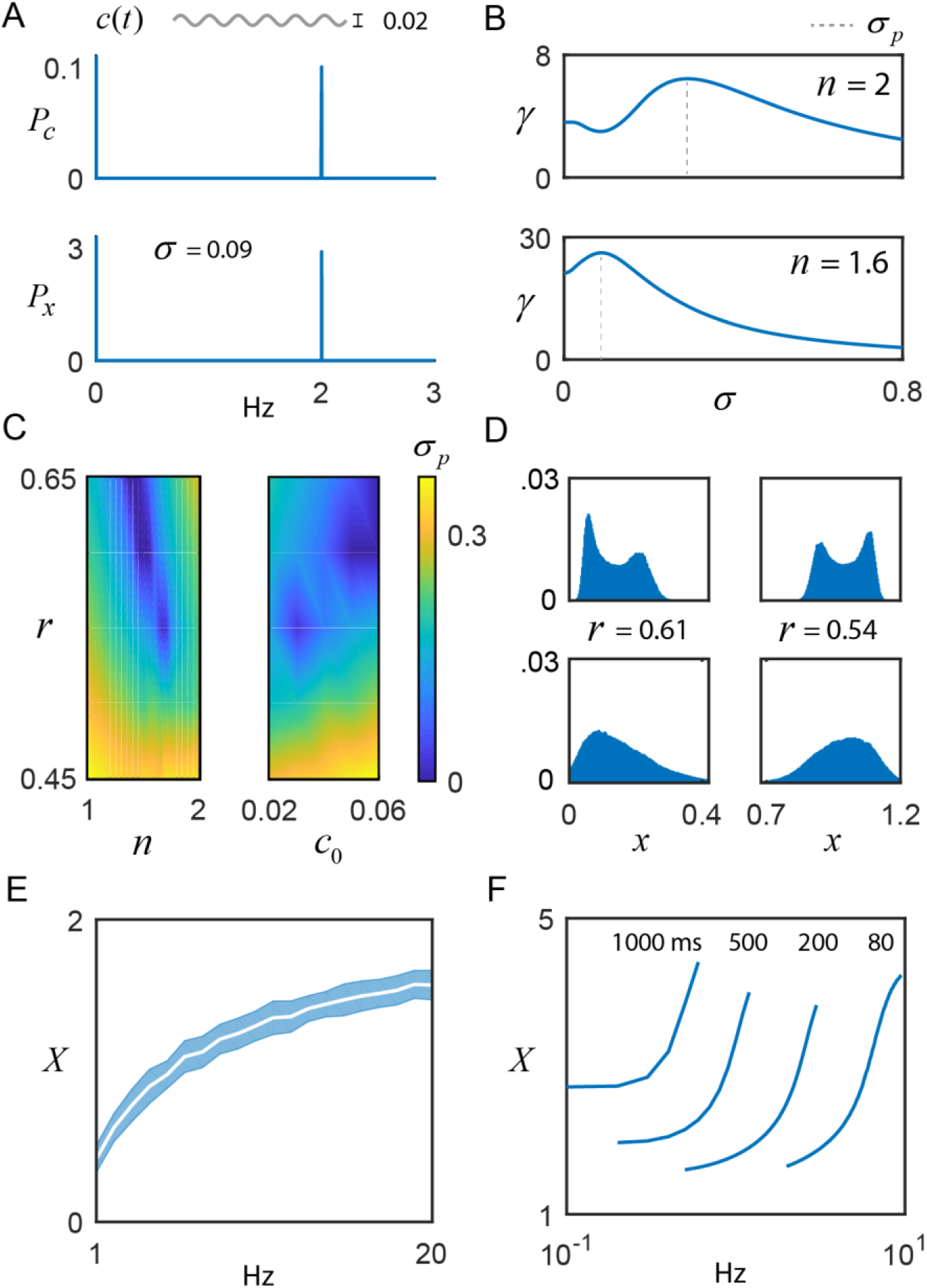
Frequency coding with noisy switches. **A)** The switch model driven by a weak sinusoidal signal, *c*(*t*) = *c*_0_ + α sin(2*φπt*), with *c*_0_ = 0.04, *α* = 0.02, *φ* = 2 Hz, and additive noise, *η*(*t*), whose intensity is scaled by the parameter *σ* and evolves according to *τ_η_* = 0.01. The switch amplifies the frequency content of the input, as shown by its power spectrum *P_x_* relative to the signal’s, *P_c_*. **B)** *Top:* For *n* = 2, the ratio of switch power to signal power at *φ* is plotted as a function of the noise intensity *σ*, achieving a maximum at 0.29, that is, the switch displays stochastic resonance (SR). The value of *σ* that promotes optimal frequency transfer is denoted by *σ_p_*. *Bottom:* For *n* = 1.6, there is substantially larger gain in the SR effect, and *σ_p_* shifts to 0.09. **C)** *σ_p_* is plotted as a function of (*n, r*) and (*c*_0_, *r*), illustrating the presence or absence of SR. **D)** For *n* = 1.6, stochastic switch simulations produce bimodal (e.g., *σ* = 0.01) or unimodal (e.g., *σ* = 0.035) activation around the low state (left column; *r* = 0.61, *c_0_* = 0.04) and the high state (right column; *r* = 0.54, *c_0_* = 0.04) (Box 1). Within each sub-state region, the input is uniquely encoded. **E)** When driven by multiple realizations of Poisson input, the autonomously activated switch (*X*), generated by Equations 2 and 3, logarithmically compresses the mean input frequency (shading: +/- standard deviation) over the course of stimulation, while instantaneous switch activity tracks the different instantaneous input frequencies as in Fig. 2A (parameters, *n* = 1.6, *r* = 0.61 and *c*_0_ = 0.04). F) As a model validation, the pulse duration (in ms) and frequency experiments of De Koninck and Schulman (1998) were simulated (*n* = 1.6, *r* = 0.61 and *T* = 0.4), qualitatively capturing their results, as well as the model results of Dupont et al. (2003). The reader should note the ambiguity in autonomously activated (long timescale) switch activity, based on input duration and frequency [38].

The model results of Kang et al. [47] depend on a full complement of parameters, which begs the question of whether stochastic resonance is a generic feature of the model switch or whether the effect is only significant for a certain range of the parameters. The dimensional reduction of the switch model performed here allows this question to be easily addressed as a function of the parameters *c*_0_, *r* and *n*. Figure 3C shows that the parameter *r* has significant influence over the value of *σ* that produces optimal spectral amplification and that, for some combinations of *c*_0_, *r* and *n*, the stochastic resonance effect disappears completely. The presence or absence of stochastic resonance may prove useful for deducing parameter ranges of molecular switches *in vitro* and *in vivo*. Furthermore, these noise fluctuations may generate unimodal (e.g., *σ* = 0.035) or bimodal (e.g., *σ* = 0.01) distributions of switch activation (Fig. 3D, *n* = 1.6), which provides another experimentally testable prediction for αCaMKII, given that the switch state could control neurotransmitter release (see Box 1) and thus explain multimodal distributions of excitatory postsynaptic potential amplitudes [60]. The occupation of the low state (Fig. 3D, left) versus the high state (Fig. 3D, right) depends on the level of negative regulation *r* and whether *c*_0_ can support hysteresis: the parameter choices for the left column of Fig. 3D do not support bistability (*r* = 0.61, *c_0_* = 0.04) and the switch fluctuates around the low activation state. The right column of Fig. 3D does support bistability (*r* = 0.54, *cα* = 0.04) and input activity quickly drives high switch activation, while hysteresis ensures the switch stays within this state. Stochastic simulations for Figure 3D were performed by including additive Ornstein-Uhlenbeck noise, as described above. Further detail can be found in the Methods section.

### Bridging short term dynamics with long timescale switch activation

A potential caveat of the bistable switch model is that, even in the high activation state, the population of phosphorylated units (*x*) are still subject to the phosphatase activity (*r*). Equation 2 places difficult constraints on cells for long-timescale activation: if *c*_0_ and *r* are not controlled carefully, the high activation state can be lost. Although high activation levels may only be short lived *in vivo*, it is important to establish a potential connection between the current model and theories of long timescale activation (Box 1; Introduction). Equation 2 effectively represents all of the phosphorylated subunits in a population of CaMKII molecules (each having twelve phosphorylation sites). When one of these dodecamers becomes fully phosphorylated, it could effectively become impervious to negative regulation by the phosphatases, since any cleaved subunit could immediately be re-phosphorylated by its neighbouring subunits and the enzyme can be shielded by its interactions with downstream targets (e.g., an NMDA receptor subunit) [14, 25]. Until now, the work presented here has ignored this potentially important feature of CaMKII, since the actual biological relevance of autonomous activation is still in question (Box 1). Therefore, to connect the short term dynamics to long timescales, we introduce a new variable (*X*) to represent the level of autonomously activated switch that might persist after the stimulus has been removed, even when Ca^2^+ levels drop below the leftmost critical value *c_c_* that supports hysteresis (Fig. 1Bi). *X* is calculated from Equation 2 by using Equation 3, explained below.

Motivated by the work of Pinto et al. [38] (Box 1), let us assume that the total amount of autonomously activated switch (*X*) is simply proportional to the average amount of Ca^2^+ input, which is determined by pulse amplitude, duration and frequency. As seen in Fig. 2, this value is reflected by the amount of activated switch *x*(*t*) over the duration of the stimulus, Δ*t*. Therefore, let *X* be the temporal average of *x* (*t*)

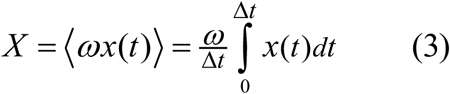

The biological interpretation is as follows: at a given moment in time there is some likelihood for individual dodecamers to transition to the fully autonomous, phopsphyorylated switch state. These autonomous elements accumulate over time. For simplicity, a basal rate of transition of a given molecule to the fully autonomous state, is assumed.

Figure 3E shows the amount of autnomously activated noisy switch, averaged over repeated realizations of Poisson input for a range of mean frequencies, as determined by Equation 2 with additive noise and Equation 3 (*n =* 1.6, *r =* 0.61, *c*_0_ = 0.04 and *σ* = 0.09). Note that the amount of autonomously activated switch performs a logarithmic compression of the mean frequency over the course of stimulation (Equation 2 and 3), while the instantaneous levels of phosphorylated switch (Equation 2) track the instantaneous input frequencies throughout each trial (e.g., Fig. 2A). Changing the parameters does not qualitatively affect the results. The reader should note that the noise in the model introduces some variability into the size of the Ca^2^+ pulse, but the averaged switch response still produces a monotonic encoding of frequency. However, if the pulse size or duration is consistently different, then ambiguity will certainly be introduced into the putative frequency code, as illustrated in Figure 3F (below), which is a replication of the original experimental data [11] that orignally prompted this criticism [38].

As a validation of the model’s ability to produce CaMKII-like behaviour over long timescales, the essence of De Koninck and Schulman’s experimental results ([11]; see Fig. 4 within) and the model of Dupont et al. ([17]; Fig. 3E within) are both captured qualitatively by Equations 2 and 3 (Fig. 3F). Note that this result was generated using Equation 2 and 3, but does not include Ornstein-Uhlenbeck noise given the synthetic and controlled nature of the original experiment [11]. The timescale factor *T* was set on the order of 10^−1^ seconds, which may reflect altered kinetics under the artificial conditions of the experiment, or the need for further refinement of the model presented here. For instance, the proportion *ω* is expected to grow larger as more of the subunit population becomes phosphorylated and cooperative activation grows stronger [61, 62], leading to an increased likelihood for individual dodecamers to transition to the fully autonomous state. This is expected to improve the reproduction of De Koninck and Schulman’s results by flattening the curves at lower frequencies and steepening them at higher frequencies [11]. Future work should seek to determine ω(*x*), with the hopes of identifying reduced representations of strongly nonlinear CaMKII activation.

## Discussion

A main goal of this study was to extend the frequency coding idea of De Koninck and Schulman [11] in a generic switch model that captures the qualitative behaviour of CaMKII, but focuses on fast timescale dynamics instead of slow timescales (Box 1). The model presented here may help to conceptually reconcile contradictory perspectives of CaMKII function [11, 38] and suggests dual streams of information transfer that are temporally multiplexed: over short timescales, where the size and duration of the Ca^2^+ pulse are more stable [39], the molecular switch might act as an encoder of instantaneous frequency information (e.g. Fig. 2A) and function to bidirectionally regulate transmitter release at synapses through a combination of enzymatic and non-enzymatic activity (summarized in Box 1). Over longer timescales, the model switch integrates overall signal intensity [15], which could dictate long term changes in synaptic strength and is dependent on multiple factors such as slow Ca^2+^- induced Ca^2^+ release (affecting *c*_0_) [31, 63], the size of the Ca^2^+ pulse, its duration and the mean frequency of stimulation (Fig. 3E). Importantly, the work presented here provides some testable predictions for synaptic physiologists: establishing the presence of both bimodal and unimodal synaptic release that depends on αCaMKII and noise, as well as characterizing the hypothesized real-time modulation of release probability at central synapses by αCaMKII in response to natural, aperiodic stimulation patterns (particularly bursting). Of particular interest is the putative role of αCaMKII in the regulation of synchronous discharge probability and duration, as well as the propagation of CA3 oscillations into the CA1 area [32].

Bistable molecular switches, such as CaMKII, are a conserved feature of cell signalling networks and generate combinatorial power in their collective action [64–66]. Due to their complex kinetics, CaMKII models are typically formulated by detailed parameterized systems of differential equations that are not readily amenable to deeper mathematical analysis. Much in the way that the leaky-integrate and fire model has been a successful abstraction of neuronal spiking activity, providing a trade-off between performance and a reduced description that facilitates network studies [22, 67], it’s proposed that simple models like Equation 2 can capture the core essence of molecular switches. This idea is supported by the inclusion of Equation 2 in a phenomenological model of feedback-driven synaptic plasticity [48]. The relative simplicity of the switch model and its application to diverse signalling pathways (e.g., MAPKs) make it a useful framework for further theoretical and experimental investigations into signalling networks, synaptic plasticity and cellular computation.

## Acknowledgements

I would like to thank Lin Wang for useful discussions and guidance during the early stages of the project, as well as Leonard Maler, Richard Naud and Jean-Claude Béïque for reading the manuscript and providing valuable comments. This work was funded by training grants awarded to S.E.C. from both the National Sciences and Engineering Research Council of Canada and the Canadian Institute of Health Research.

## Methods

### Bifurcation Analysis

The first step of the bifurcation analysis is to find the equilibrium points. Setting *n* = 2, we rewrite Equation 1 as,

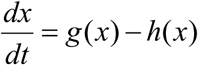
where 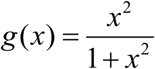 and *h*(*x*) = *rx − c*. The fixed points occur when *g*(*x*) − *h*(*x*) *=* 0, which amounts to finding the solutions of the polynomial −*rx*^3^ + (*c* + 1)*x*^2^ − *rx + c =* 0. First, fix *c* and examine the effects of varying *r*. When *c =* 0, *x* = 0 is a fixed point, and, for a particular range of *r*, there exists two other positive valued fixed points, given by the roots of *−rx^2^ + × − r =* 0. The critical value of the parameter *r*, denoted by *r_c_* is found by setting *g*(*x*) = *h*(*x*) and *g*′(*x*) = *h*′(*x*), which, when solved, gives 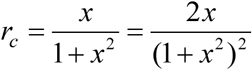. Three values of x satisfy this equality: −1, 0 and 1. Since we are not considering negative values of *x*, we have two critical points, *r_c_* = 0 and 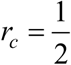. Therefore, when *c* = 0, the system is bistable for 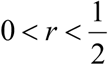. For *c* > 0, *r* can be larger than 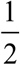 while still preserving bistability (as in Fig. 1A). We know *r_c_* occurs when *h*(*x*) = *g*(*x*) and *h*′(*x*) = *g*′(*x*); therefore, when *h*(*x*) > *g*(*x*) we lose a fixed point through a saddle node bifurcation. For *x* > 0, the maximum of *g*(*x*) is found at 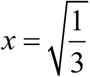 which gives 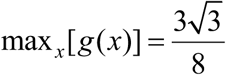. Therefore, when 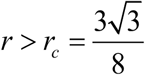 only one fixed point exists.

Now, we are interested in fixing *r* and examining the effects of varying *c*. To find *c_c_* we set *g*(*x*) = *h*(*x*) and *g*′(*x*) = *h*′(*x*), which gives 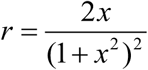 and 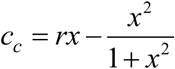. Substituting the first expression into the second, we get 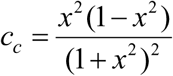. We differentiate with respect to *x* in order to locate the maximum value for 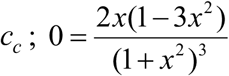. This gives *x* = 0 and 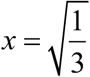, which corresponds to *c_c_* = 0 and 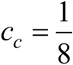. When *c* > *c_c_*, only one fixed point exists for all values of *r*. For a fixed value of *r* that supports bistability, as *c* increases from 0 and crosses a critical value (*c_c_*), the fixed point *x*\* will jump up to the high amplitude branch. If *c* is now decreased, the fixed point remains on the high amplitude branch even as *c* becomes smaller than the corresponding *c_c_*. This hysteresis effect permits switch activation to remain as the transient Ca^2+^ signal subsides, consistent with the findings from synaptic plasticity experiments (Box 1). Using the expressions derived for the critical values of *r_c_* and *c_c_*, we plot them parametrically as functions of *x* (Fig. 1Biii). Saddle node bifurcations occur all along the boundary of these curves, it is here we find the values of *r* and *c* for which only two fixed points occur. Crossing each branch results in a pairwise collision and disappearance of two fixed points. Note where the bifurcation curves meet tangentially, 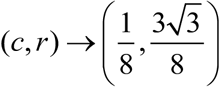, here we observe a co-dimension two bifurcation; beyond this point there is only one fixed point and the distinction between low and high activation states is blurred (Fig. 1Biii).

### Computational Specifications and Miscellaneous Details

Simulations were solved using the 4^th^ order Runge-Kutta method, with the exception of the Ornstein-Uhlenbeck noise, which was solved using the stochastic Euler method (time step of 1 ms in all cases). All simulations were performed using custom code, available upon request to the author, and were implemented on a Linux machine running Ubuntu 16.04 with an Intel core i7–6700 CPU, 3.4 GHz processing speed, and 62 GB of RAM.

Pulse train sequences {*t_i_*} were convolved with the filter 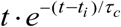, whose decay constant *τ_c_* was set to 30 ms, reflecting an accommodation of both pre- and post-synaptic calcium decay values from the literature that range from 15–43 ms [51–53]. The resulting input signal was normalized to the maximum value and then scaled by Δ*c*. The decay value is closely related to the input frequencies typical of a given synapse and the definition of what constitutes a high frequency event in the system, since for events occurring faster than the decay, Ca^2+^ accumulates quickly, driving the switch into the upstate. The putative burst detector will work for different *τ_c_*, but may require a different set of corresponding switch parameters, range of stimulation frequencies and pulse amplitudes.

Histogram bin sizes for Fig. 3D were set using the Freedman-Diaconis method [68].

